# KIF3C Regulates Bergmann Glia Density and Patterning during Cerebellar Development

**DOI:** 10.1101/2024.12.02.626415

**Authors:** Bridget Waas, Benjamin L. Allen

## Abstract

Kinesin-2 motors have been demonstrated to regulate Hedgehog (HH) signaling, particularly within the developing cerebellum (Spassky et al., 2008; Waas et al., 2024). KIF3A/KIF3B are required for ciliogenesis in mice, and loss of this motor complex results in cerebellar hypoplasia due to lack of mitogenic response of HH-responsive cerebellar granule neural progenitors (CGNPs). Homodimeric KIF17 regulates HH-dependent CGNP proliferation through regulating levels of SHH and GLI3 but is not required for ciliogenesis in the developing cerebellum. Germline deletion of *Kif3c* has previously been reported to have no effect on embryonic development or gross cerebellar morphology in adult mice (Yang et al., 2001). Examination of these mice during neonatal development reveal cerebellar hypoplasia due to reduced CGNP proliferation in *Kif3c* mutant mice. While decreased CGNP proliferation is often associated with reduced Hedgehog (HH) signaling, we did not detect a change in the levels of HH signaling in *Kif3c^-/-^* cerebella. However, we observed Bergmann glia density and localization is altered in *Kif3c* mutants, a phenotype consistent with disrupted Notch signaling. Further, we detected reduced expression of *Hes1* in *Kif3c^-/-^* cerebella. Altogether, these data demonstrate KIF3C is required for proper cerebellar development but in a HH-independent manner.

**Highlights:**

- *Kif3c* is widely expressed in developing cerebellum
- KIF3C regulates cerebellar size, but in a HH-independent manner
- *Kif3c^-/-^* mice have increased BG density and abnormal spatial patterning

## Introduction

The postnatal expansion of the cerebellum is dependent on several mitogenic pathways. Sonic Hedgehog, SHH, induces proliferation of cerebellar granule neurons (CGNPs), which give rise to cerebellar granule neurons, the most abundant type of neuron in the brain (Dahmane and Ruiz i Altaba, 1999; Wechsler-Reya and Scott, 1999). Deletion of *Shh* results in reduced proliferation and cerebellar hypoplasia (Lewis et al., 2004). Additionally, loss of other essential HH pathway components, such as *Gli2*, *Gas1* or *Boc*, negatively impact CGNP proliferation and cerebellar size. Another signaling pathway required for proper cerebellar development is Notch signaling (Adachi et al., 2021; Hiraoka et al., 2013; Komine et al., 2007; Solecki et al., 2001; Weller et al., 2006). Stimulation of the Notch pathway with JAG1 induces proliferation and inhibits differentiation of CGNPs (Solecki et al., 2001). Deletion of Notch pathway components, such as *Jag1*, *Notch1*, *Notch2* or *Dll-1*, results in CGNP migration defects and abnormal localized Bergmann glia, which undergo apoptosis (Adachi et al., 2021; Hiraoka et al., 2013; Komine et al., 2007; Weller et al., 2006).

Kinesin-2 motors have been demonstrated to regulate Hedgehog signaling (Huangfu et al., 2003; Spassky et al., 2008). The kinesin-2 family consists of three motor complexes: heterodimeric KIF3A/KIF3B, homodimeric KIF17 and heterodimeric KIF3A/KIF3C [reviewed in (Hirokawa et al., 2009)]. Heterodimeric KIF3A/KIF3B is well known for its role in ciliogenesis and HH signaling during embryogenesis (Engelke et al., 2019; Huangfu et al., 2003; Nonaka et al., 1998; Takeda et al., 1999). Further, KIF3A/KIF3B physically interacts and regulates with HH transcriptional effectors, GLI transcription factors (Carpenter et al., 2015). Homodimeric KIF17 has been found to regulate HH signaling at the level of SHH ligand and the level of GLIs in the developing cerebellum (Waas et al., 2024).

The remaining motor complex KIF3A/KIF3C has not been investigated for a role in HH signaling or cerebellar development. Unlike its other family members, KIF3C loss does not result in ciliary phenotypes, and *Kif3c* deletion is well tolerated across multiple model organisms (Jimeno et al., 2006; Yang et al., 2001; Zhao et al., 2012). However, *Kif3c* mutant mice display defects in neuronal regeneration (Gumy et al., 2013). Specifically, growth cones of *Kif3c^-/-^* neurons displayed stable, overgrown, and looped microtubules that impair regeneration (Gumy et al., 2013). Given the roles of other kinesin-2 motors in HH-dependent cerebellar development, we investigated the contribution of the kinesin-2 motor, KIF3A/KIF3C, in HH signaling within the developing cerebellum.

Here we find that *Kif3c* is widely expressed in the developing cerebellum, specifically within CGNPs, Bergmann glia, Purkinje cells and CGNs. Germline deletion of *Kif3c* results in cerebellar hypoplasia and reduced body size postnatally. We observe reduced *in vivo* proliferation of CGNPs in *Kif3c^-/-^*mice, but unexpectedly, HH signaling is not affected. Increased Bergmann glia density and disrupted spatially patterning was observed in *Kif3c* mutant cerebella, and reduced expression of *Hes1* suggests KIF3C is required for proper cerebellar development, potentially through regulating Notch signaling. Altogether, these data demonstrate KIF3C is required cerebellar development, but in a HH-independent manner.

## Methods

### Animal models

*Kif3c* germline mutant mice have been previously described (Yang et al., 2001). Mice were maintained on congenic C57BL/6J background. All animal procedures were reviews and approved by the Institutional Animal Care and Use Committee (IACUC) at the University of Michigan, USA. Experiments performed in this paper were completed with littermate controls.

### RT-qPCR

Cerebella were dissected in 1X PBS, and RNA was isolated using a PureLink RNA Mini Kit (ThermoFisher Scientific, 12183025). Following isolation, 2 µg of RNA were used to generate cDNA libraries using a High-Capacity cDNA reverse transcription kit (Applied Biosystems, 4368814). RT-qPCR was performed using PowerUP SYBR Green Master Mix (Applied Biosystems, A25742) in a QuantStudio 3 Real-Time PCR System (Applied Biosystems). Primers used in this paper can be found in Table 1. Gene expression was normalized to *Gapdh,* and relative expression analyses were performed using the 2(-ddCT) method. For RT-qPCR analysis, biological replicates were analyzed in triplicate.

### Wholemount digoxigenin in situ hybridization

Wholemount digoxigenin in situ hybridization was performed as previously described (Allen et al., 2011; Wilkinson, 1992). First, cerebella were dissected in 1X PBS (pH 7.4), cut in half with a razor and fixed for 24 h with 4% paraformaldehyde at 4°C on rocking platform. The *Kif3c* probe was designed to bind to the end of the mRNA transcript: (GCTGCTCCACTGGACTGAATGGCGGAGCCTTGCGGCTGCCTGCCCTTCAAAGGGATCCCAGGTTTCTGTCAGAACCCTGTGATTGACACTCAGGATTCAAATCAGAGGAATGGCTTTCTCTGGAACAGGAGCTGTGTGTAGAAATCTCCTGATGTGAACTGGGCATTGAGGGACCTCCCCCTGAGCTCTCTGTCATTTGTAGATGAAGCTGCATGAGTCACCCCATTCATCACTTGGACACACTGACTCCACATTGTCTGGTCCACTACCCTCACAGTCTTATAGCACAATACACCCCACTTCAGCACCGCAGCCAAAGGCTGGGCCCAAGGTGTGGTCAGAAGAGGTGCTCCTGCCTGTGGTATTATATGTGTGTGTTTATGTGTGTGTTTATGTTCACCTGTACAGGGGGCACTACACTCAATGTAAGATACCCTGGAGACAGGACTCCTGGAGGTGGCTGGATCTCAGTCTCTGTCTCTCTCCTTTTTCTTTTACTGTATCACACATTTGATTGACAAAGTACGGGCCTTAATTAGGATCAAATTTCTATGTCTGTTGCTATGGCCTTTAATTAAAGTTACACAAAGTGGCCCATTCTTGTCACTCTATACATATGGGACATATGTATATCTAGGACATATGTAATATATAAATATATAAATATATATAAAGCATTAACCTCTGCCCCC). Probe hybridization was performed with the *Kif3c* digoxigenin probe at a concentration of 1 ng/µl overnight at 70°C. The samples were incubated in AP-conjugated anti-DIG antibody (Table 2). AP-anti-DIG was visualized with BM Purple (Roche, 11442074001), and signal was developed for 4 h at 37°C. After the signal was developed, development was stopped with 3 x 5 min washes with 1X PBS (pH 4.5). Cerebella were post-fixed in 4% PFA + 0.2% glutaraldehyde for 30 min, then washed 3 x 5 min in 1X PBS (pH 7.4). Cerebella were photographed using a Nikon SMZ1500 microscope and stored in 1X PBS (pH 7.4).

### Fluorescent in situ hybridization

Cerebella were dissected in 1X PBS (pH 7.4) and cut in half using a razor. Cerebella were fixed with 10% neutral buffered formalin (Fisher, 245-685) on a rocking platform at room temperature for 24 h. Following fixation, cerebella were washed 3 x 5 min with 1X PBST^X^ on a rocking platform and cryoprotected overnight in 1X PBS + 30% sucrose on a rocking platform. Cerebella were then washed 3 x 1 h with 50% OCT compound before embedding in 100% OCT. Sections were collected on a Leica CM1950 cryostat at 12 µm thickness. Slides were processed using RNAscope Multiplex Fluorescent Detection kit (ACD, 323110) using a protocol adapted from (Holloway et al., 2021). Prior to probe hybridization, samples underwent antigen retrieval for 15 minutes and treated with Protease Plus (ACD, 322381) for 5 minutes. Probes used in this paper were a custom probe for *Mm-Kif3c*, designed to bind to exon 1, and *Mm-Gli1* (ACD, 311001). After probe detection, slides were subsequently stained using the below-described section immunofluorescence protocol.

### Section Immunofluorescence

Section immunofluorescence was performed as described in (Allen et al., 2011). Briefly, cerebella were dissected in 1X PBS (pH 7.4) and cut in half using a razor. Cerebella were fixed with 4% paraformaldehyde (Electron Microscopy Sciences) for 1 h on ice. Following fixation, cerebella were washed 3 x 5 min with 1X PBS (pH 7.4) on a rocking platform and cryoprotected overnight in 1X PBS + 30% sucrose on a rocking platform. Then, cerebella were washed 3 x 1 h in 50% OCT (Fisher Scientific, 23-730-571) before embedding in 100% OCT. Sections were collected on a Leica CM1950 cryostat at 12 µm thickness for all experiments. Slides were then washed 3 x 5 min with 1X PBS (pH 7.4). For mouse primary antibodies, citric acid antigen retrieval (10 mM citric acid + 0.5% Tween-20, pH 6.0) at 92°C for 10 min was performed prior to primary antibody incubation. Primary antibodies were diluted in blocking buffer (3% bovine serum albumin, 1% heat-inactivated sheep serum, 0.1% Triton X-100) and incubated overnight at 4°C in a humidified chamber. After primary antibody incubation, slides were washed 3 x 10 min with 1X PBST^X^ (1X PBS + 0.1% Triton X-100, pH 7.4). Secondary antibodies were diluted in blocking buffer and incubated for 1 h at room temperature, followed by 3 x 5 min 1X PBST^X^ washes. Nuclei were labeled using DAPI (0.5 µg/mL in blocking buffer) for 10 min and washed twice with 1X PBS. Coverslips were mounted using Immu-mount aqueous mounting medium (Thermo Fisher Scientific, 9990412). Images were taken on a Leica SP5X upright confocal (2 photon). A list of all the primary and secondary antibodies and their working concentrations is provided in Table 2. For analysis, we examined mid-sagittal cerebellar sections, where lobes I-III were considered anterior, while lobes VI-VIII were considered posterior.

### Weight analyses

For weight measurements, the date litters were born were noted as postnatal day 0 and were dissected on postnatal day 10. Pups were first weighed and then placed on ice briefly before decapitation. The cortices and cerebella were dissected in 1X PBS (pH 7.4). To weigh cortices and cerebella, a specimen jar was first filled with PBS on an analytical scale. The tissue was transferred with forceps to the specimen jar, and its weight was recorded. Genotyping samples were taken after dissection, allowing the weights to be recorded without prior knowledge of the genotype.

### EGL and PC Dendrite quantitation

To measure the thickness of the external granule layer (EGL) and PC dendrite length, ImageJ software was utilized. Images were first blinded before measuring. For EGL thickness, the area was divided by the length of the EGL. For PC dendrite length, measurements were taken just below the bottommost nuclei in the EGL to the center of Purkinje cell nuclei within the molecular layer. For each animal, at least three images were acquired in the posterior lobes and an additional three images in the anterior lobes. For analysis, we examined mid-sagittal cerebellar sections, where lobes I-III were considered anterior, while lobes VI-VIII were considered posterior

### EdU incorporation assay

On postnatal day 9, pups were intraperitoneally injected with 100mg/kg of EdU (Invitrogen, A10044), dissolved in 1X PBS (pH 7.4). 24 h later, cerebella were dissected and processed for section immunofluorescence as described above. Prior to primary antibody incubation, EdU incorporation was visualized with an azide staining solution [100 mM Tris HCl (pH 8.3), 0.5 mM CuSO4, 50 mM ascorbic acid, 50 µM Alexa Fluor 555 Azide, Triethylammonium Salt (Thermo Fisher Scientific, A20012)] for 30 min at room temperature. Sections were then washed 3 x 10 min in PBST^X^ (1x PBS + 0.1% Triton X-100, pH 7.4), followed by immunofluorescence staining as described above.

### Image quantitation

To quantify fluorescent *Gli1* fluorescence, ImageJ software was used to measure the integrated density fluorescent signal contained to either the external granule layer (EGL, CGNPs) or lower molecular layer and inner granule layer (IGL, Bergmann glia and CGNs). The signal was then divided by the number of HH-responsive cells in each layer (number of PAX6^+^ cells in the EGL; number of PAX6^+^ and SOX2^+^ cells in the molecular layer and IGL). At least three images were analyzed per region per mouse. For all image analyses, images were blinded.

### Quantitation and statistical analysis

All the data are mean ± s.d. All statistical analyses were performed using GraphPad Prism (www.graphpad.com). Statistical significance was determined by using a two-tailed Student’s t-test. For all the experimental analyses, a minimum of three mice of each genotype were analyzed, each n represents a mouse. For in vitro experiments, a minimum of three biological replicates were analyzed, each n represents a biological replicate. All the statistical details (statistical test used, adjusted P-value, statistical significance and exact value of each n) for each experiment are specified in the figure legends.

## Results

### Kif3c is expressed in the developing cerebella and is required for proper cerebellar development

To investigate if *Kif3c* is required for cerebellar development, *Kif3c^-/-^* mice were acquired on a C57BL/6J genetic background with deletion of exon 1 (Figure S1A). Consistent with the literature and similar to *Kif17*, *Kif3c^-/-^* mice are viable and fertile (Yang et al., 2001). To determine if *Kif3c* is expressed in the developing cerebellum, RT-qPCR was performed on postnatal day 10 (P10) *Kif3c^+/+^*and *Kif3c^-/-^* cerebella (Figure S1B), where we detected expression in *Kif3c^+/+^* cerebella but not in *Kif3c^-/-^* cerebella. Whole-mount *in situ* hybridization of *Kif3c* was performed in P10 *Kif3c^+/+^*and *Kif3c^-/-^* cerebella (Figure S1C-D). Distinct from *Kif17*, *Kif3c* appears to be uniformly expressed across the lobes of the cerebellum and within all cell layers of the cerebellum. To determine which cell populations express *Kif3c*, fluorescence *in situ* hybridization was performed in anterior lobes and posterior lobes of *Kif3c^+/-^* (Figure 1A-B, E-P) and *Kif3c^-/-^* (Figure 1C-D, Figure S1E-P) P10 cerebella. In both anterior and posterior lobes, *Kif3c* puncta were observed in CGNPs (Figure 1E-H), surrounding Purkinje cell nuclei, Bergmann glia nuclei (Figure 1I-L) and within mature CGNs (Figure 1M-P). This data suggests *Kif3c* is expressed across multiple cell types in the developing cerebellum.

**Figure 1:**
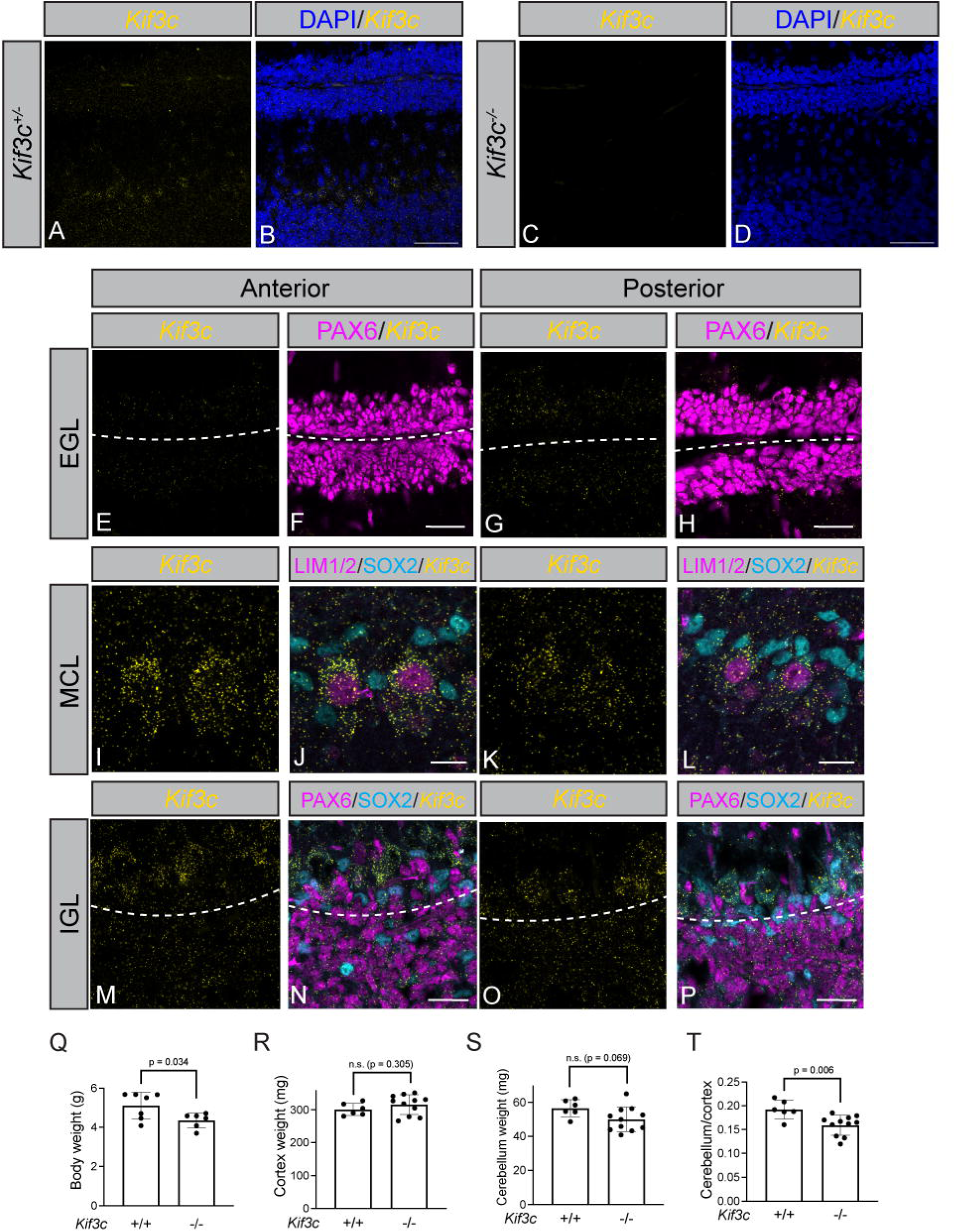
*Kif3c* is expressed in the developing cerebellum and required for proper cerebellar size. Fluorescent *in situ* detection of *Kif3c* mRNA (yellow; **A-P**) in *Kif3c^+/-^* and *Kif3c^-/-^* anterior and posterior lobes in P10 cerebella. DAPI (blue, **B, D**) to label all nuclei. Antibody detection of PAX6 (magenta) to label cerebellar granule neural progenitor nuclei. Dashed lines separate individual external granule layers. Antibody detection of LIM1/2 and SOX2 (magenta, cyan; F, H) to label Purkinje cells and Bergmann glia, respectively. Antibody detection of PAX6 and SOX2 (magenta, cyan; J, L) to label CGNs and Bergmann glia, respectively. Dashed lines separate molecular layer (MCL) and inner granule layer (IGL). Scale bars, (**B, D**), 50 µm. Scale bars (**F, H**), 15 μm. Scale bars (**J, L, N, P**), 25 μm. Quantitation of body weight (**Q**), cortex weight (**R**), cerebellar weight (**S**), and cerebellar weight normalized to cortex weight (**T**) in P10 *Kif3c^+/+^*and *Kif3c^-/-^* mice. Data are mean ± s.d. Each dot represents an individual animal. P- values were determined by a two-tailed Student’s t-test.

To investigate whether *Kif3c* deletion affected gross postnatal development, we analyzed body, cortical and cerebellar weights. We observed *Kif3c* deletion results reduced body size at during first three postnatal weeks (Figure 1Q, Figure S2A). Importantly, the difference between *Kif3c^+/+^* and *Kif3c^-/-^* body sizes is not significant by 6 weeks of age, distinct from HH loss-of- or gain-of-function animals, which maintain significant differences during adulthood (Goodrich et al., 1997; Zhang et al., 2015). We did not detect a difference in cortical weights (Figure 1R), and examination of cerebellar weights revealed a non-significant reduction in cerebellar size (Figure 1S). However, after normalizing the cerebellar weight to the cortical weight (Figure S2E), the difference in *Kif3c^-/-^*mice was significant, demonstrating the requirement for *Kif3c* in the developing cerebellum. Altogether, these data demonstrate *Kif3c* is required for proper cerebellar size and expressed in multiple cell types in the developing cerebellum.

### Kif3c deletion results in reduced CGNP proliferation in a HH-independent fashion

To investigate which cells were affected by *Kif3c* deletion in the developing cerebellum, we measured the length of Purkinje cell (PC) dendrites and thickness of the external granule layer (EGL), where CGNPs reside, were measured (Figure 2A-D). We did not detect a difference in PC dendrite length in the anterior (Figure 2A) or posterior (Figure 2C) regions with *Kif3c* deletion. However, *Kif3c^-/-^* cerebella have a significant reduction in EGL thickness in both regions (Figure 2B, D). In other studies (Izzi et al., 2011), reduced EGL thickness was associated with decreased CGNP proliferation, which was examined in *Kif3c^+/+^*and *Kif3c^-/-^* littermates (Figure 2E-T). While there was not a change in the percentage of Ki67^+^ CGNPs in the anterior lobes (Figure 2Q), we observe a significant reduction in the percentage of EdU^+^ CGNPs (Figure 2R). In the posterior lobes, we observe a reduction in the percentage of EdU^+^ and Ki67^+^ CGNPs (Figure 2S-T). Further, the expression proliferative CGNP marker, *Atoh1*, measured through RT-qPCR, is significantly reduced in *Kif3c^-/-^* P10 cerebella (Figure S2B). Collectively, these data suggest *Kif3c^-/-^* cerebellar hypoplasia is due to reduced CGNP proliferation.

**Figure 2:**
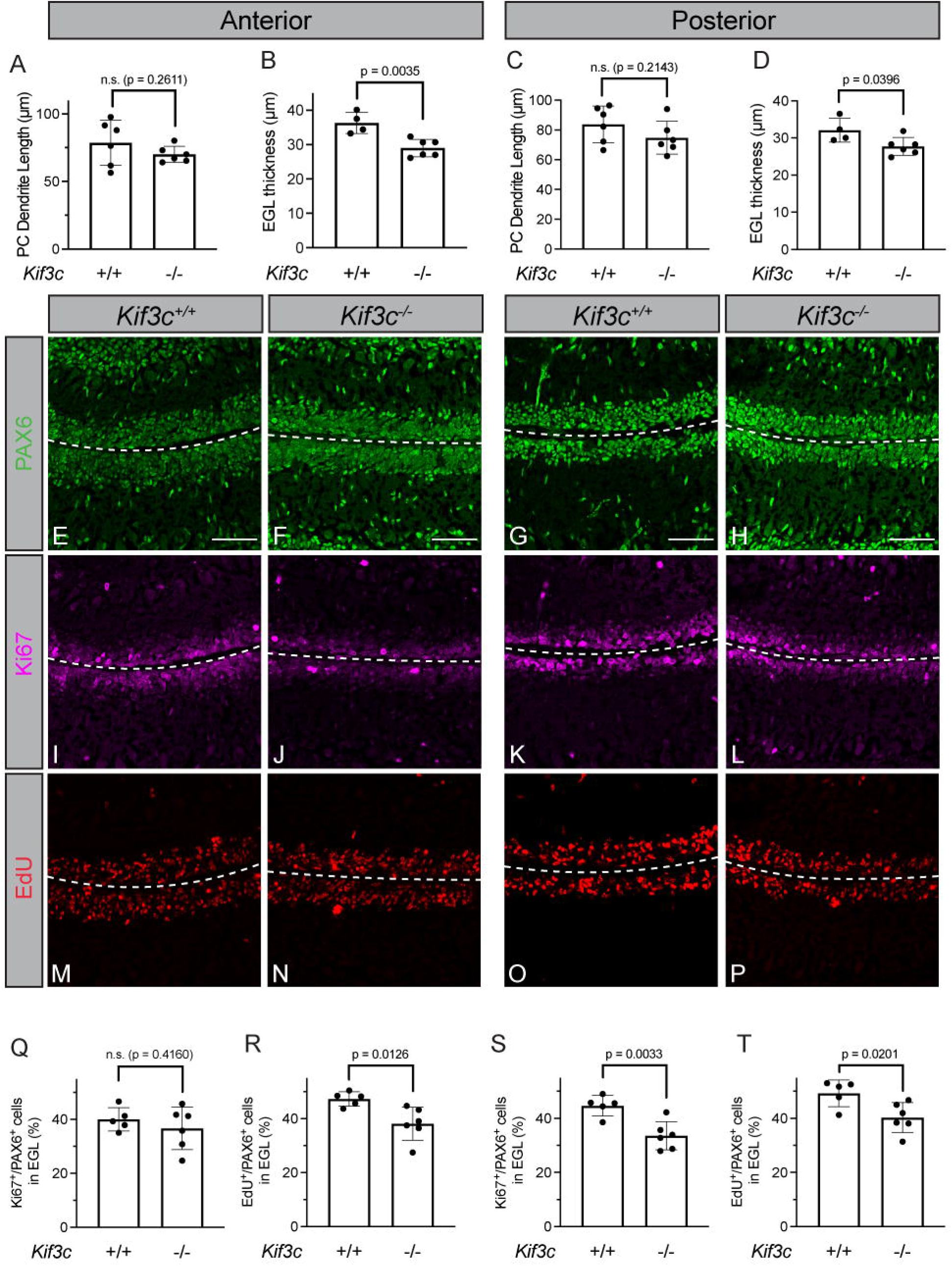
Reduced EGL thickness and CGNP proliferation in *Kif3c* mutant cerebella. Quantitation of PC dendrite length (A) and external granule layer (EGL, B) thickness in the anterior lobes of P10 *Kif3c^+/+^* and *Kif3c^-/-^*cerebella. Quantitation of PC dendrite length (C) and external granule layer (EGL, D) thickness in the posterior lobes of P10 *Kif3c^+/+^* and *Kif3c^-/-^* cerebella. Immunofluorescent analysis of CGNP proliferation of *Kif3c^+/+^* (E, F, I, J, M, N) and *Kif3c^-/-^* (**G, H, K, L, O, P**) in anterior and posterior lobes in P10 cerebella. Antibody detection of PAX6 (green; **E-H**) and Ki67 (magenta; **I-L**). Fluorescent azide detection of EdU (red; **M-P**). Scale bars (**E, G**), 50μm. Dashed line separates individual external granule layers. Quantitation of the percentage of Ki67^+^ (**Q**) and EdU^+^ (**R**) PAX6^+^ cells in the EGL in the anterior lobes of *Kif3c^+/+^*and *Kif3c^-/-^* P10 cerebella. Quantitation of the percentage of Ki67^+^ (**S**) and EdU^+^ (**T**) PAX6^+^ cells in the EGL in the posterior lobes of *Kif3c^+/+^* and *Kif3c^-/-^* P10 cerebella. Each dot represents the average of at least three images per individual animal.

As reduced levels of HH signaling are often associated with reduced CGNP proliferation and cerebellar hypoplasia (Corrales et al., 2006; Izzi et al., 2011; Lewis et al., 2004; Spassky et al., 2008), we next assessed the levels of HH signaling through fluorescence *in situ* hybridization of *Gli1* (Figure 3A-H). In the anterior and posterior lobes at P10, we do not detect any significant changes in the fluorescent intensity of *Gli1* expression in the anterior or posterior lobes (Figure S2C-D). While there is a non-significant reduction of *Gli1* fluorescence within CGNPs, when fluorescent intensity is normalized to the number of HH-responsive cells, this difference becomes less apparent (Figure 3I-J). Analysis of other HH target genes, *Ptch1*, *Ptch2*, measured by RT-qPCR, reveal no significant changes of expression (Figure 3K-L). Furthermore, expression of *Shh* is unchanged in *Kif3c^-/-^* cerebella (Figure 3M). Together, these data suggest that *Kif3c^-/-^* cerebella do not display a perturbation to HH signaling, unlike other kinesin-2 mutants.

**Figure 3:**
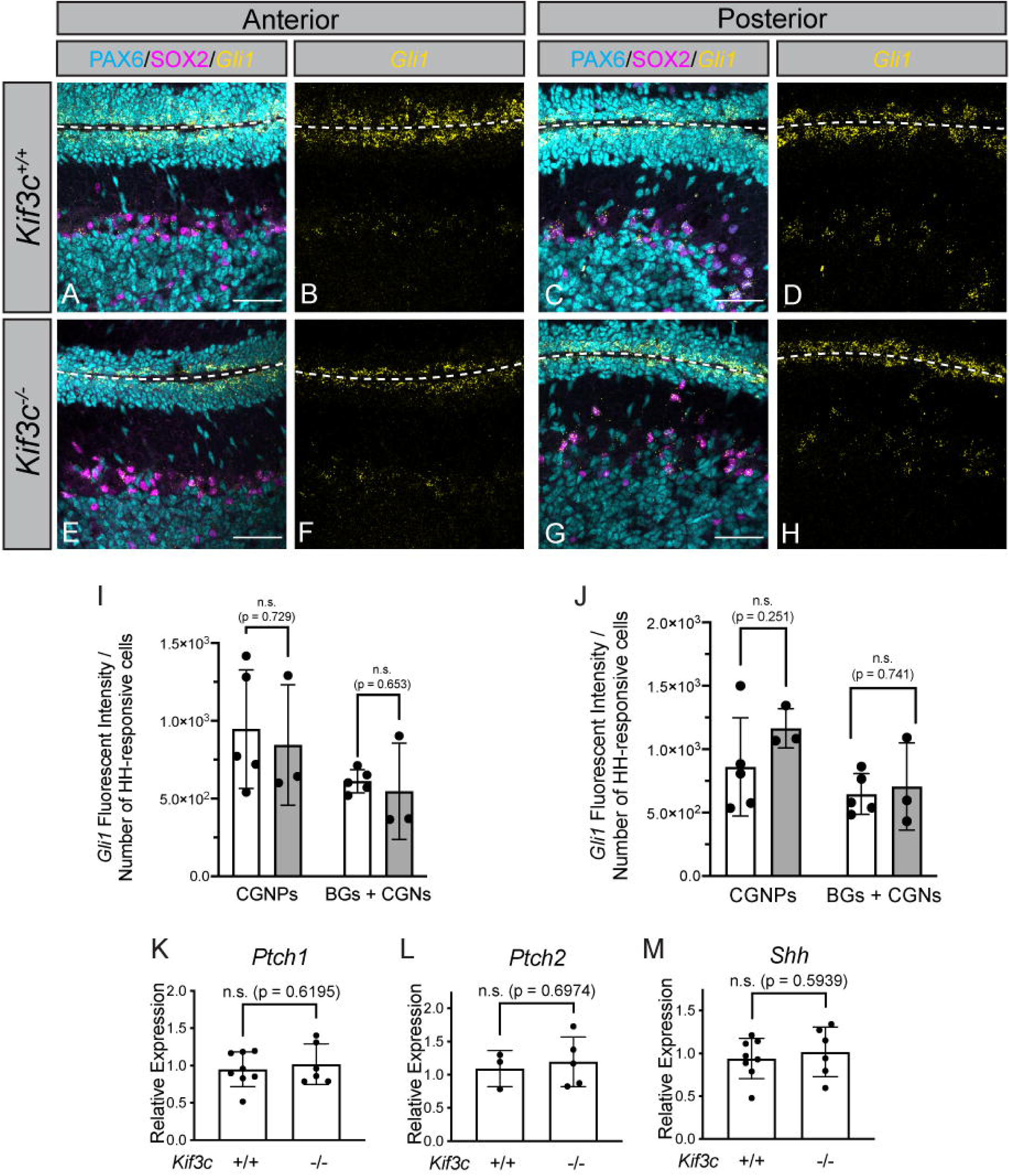
*Kif3c^-/-^*cerebella do not display perturbations in HH signaling. Fluorescent *in situ* detection of *Gli1* (yellow; **A-H**) in P10 *Kif3c^+/+^* and *Kif3c^-/-^* cerebella in the anterior (**A, B, E, F**) and posterior (**C, D, G, H**) lobes. Antibody detection (**A, C, E, G**) of PAX6 (cyan) and SOX2 (magenta) to label granule neurons and Bergmann glia, respectively. Quantitation of fluorescent intensity (integrated density) of *Gli1* puncta normalized to the number of HH-responsive cells within the EGL (CGNPs) or MCL/IGL [(Bergmann glia and cerebellar granule neurons (BGs + CGNs)] in P10 *Kif3c^+/+^*and *Kif3c^-/-^* anterior (**I**) and posterior (**J**) lobes. Scale bars (**A, C, E, G**), 50 µm. Dashed lines separate external granule layers. Each dot represents the average of at least 3 images per animal. RT-qPCR detection of *Ptch1* (**K**), *Ptch2* (**L**) and *Shh* (**M**) expression P10 *Kif3c^+/+^* and *Kif3c^-/-^* cerebella. Data are mean ± s.d. Each dot represents an individual animal. *P*- values were determined by a two-tailed Student’s *t*-test.

While examining *Gli1* expression in *Kif3c* mutant cerebella, we observed abnormal density and localization of Bergmann glia in *Kif3c^-/-^*mice (Figure 3E, G). To further investigate, we examined the density of Purkinje cells (Figure 4A, C, E, G) and Bergmann glia (Figure 4B, D, F, H) in the anterior and posterior lobes of *Kif3c^+/+^* and *Kif3c^-/-^* cerebella. We did not detect a change in the density of Purkinje cells in either the anterior (Figure 4I) or posterior (Figure 4K) lobes of *Kif3c* mutant mice. However, we did detect an increase in Bergmann glia density within both the anterior (Figure 4J) and posterior (Figure 4L) lobes of *Kif3c^-/-^* cerebella. Further, Bergmann glia spatial patterning was disturbed, particularly in the posterior lobes (Figure 4F, H, white arrowheads). Notably, some Bergmann glia resided within the EGL. This phenotype was reminiscent of previous studies of Notch signaling mutants, where there is decreased CGNP proliferation as well as abnormal Bergmann glia localization (Adachi et al., 2021; Hiraoka et al., 2013; Komine et al., 2007; Weller et al., 2006). We assessed the expression of Notch signaling components through RT-qPCR with *Kif3c* deletion (Figure 4M-N). While we did not detect a difference in Notch ligand, *Jag1* (Figure 4M), we observed a significant reduction in Notch target gene, *Hes1* (Figure 4N). Collectively, these data demonstrate *Kif3c* deletion results in reduced CGNP proliferation and abnormal Bergmann glia patterning, potentially through reduced Notch signaling.

**Figure 4:**
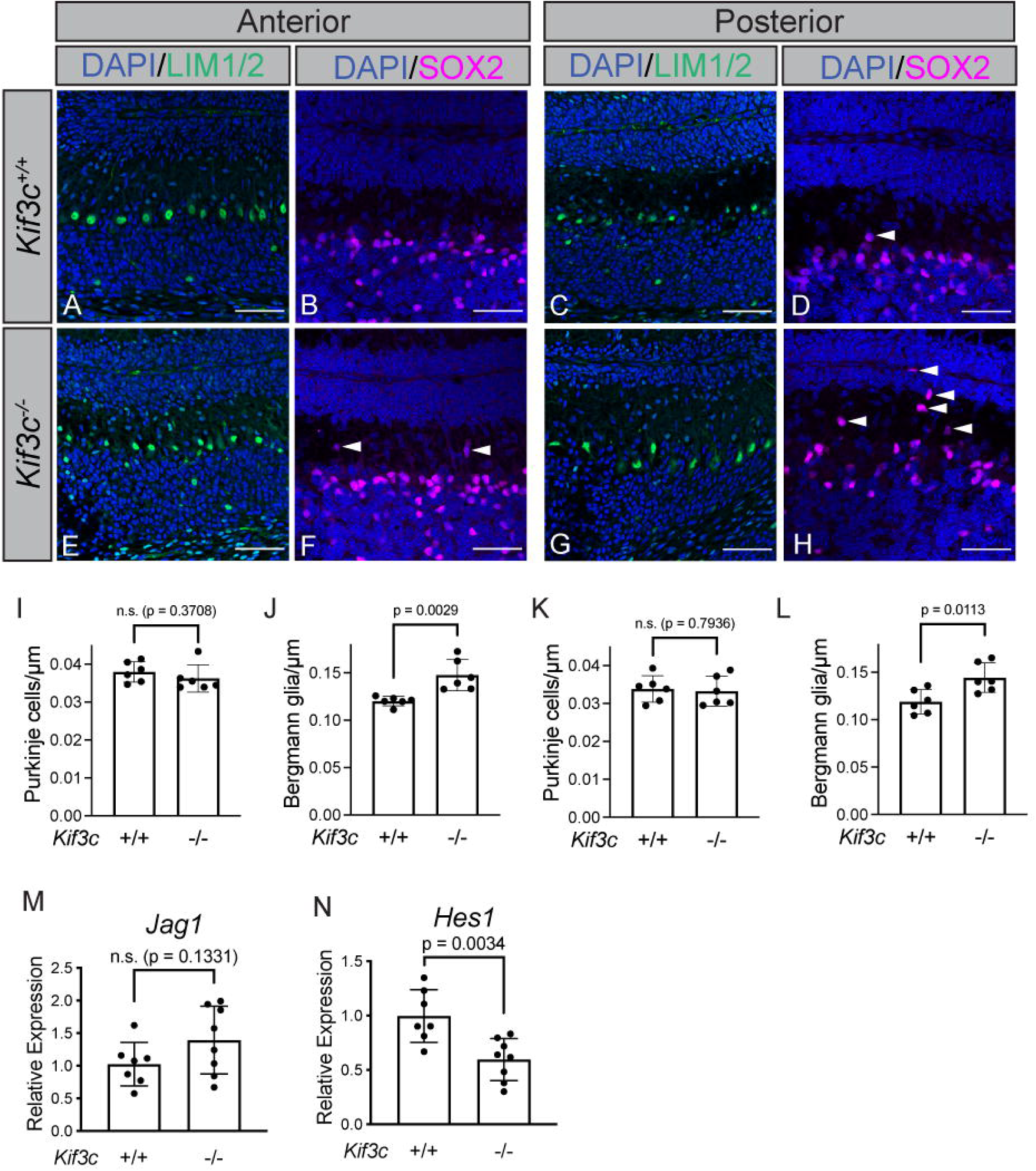
Increased density and abnormal spatial patterning of Bergmann glia in *Kif3c^-/-^* cerebella. Immunofluorescent analysis of Purkinje cell and Bergmann glia localization and density in *Kif3c^+/+^* (**A-D**) and *Kif3c^-/-^* (**E-H**) P10 cerebella in the anterior (**A, B, E, F**) and posterior (**C, D, G, H**) lobes using immunofluorescent detection of LIM1/2 (green) and SOX2 (magenta) to mark Purkinje cells and Bergmann glia, respectively. Nuclei were counterstained with DAPI (blue). Scale bars (**A-H**), 50 µm. White arrowheads denote Bergmann glia outside of the molecular layer. Quantitation of Purkinje cell (**I**) and Bergmann glia density (**J**) in anterior lobes of *Kif3c^+/+^* and *Kif3c^-/-^* P10 cerebella. Quantitation of Purkinje cell (**K**) and Bergmann glia density (**L**) in posterior lobes of *Kif3c^+/+^* and *Kif3c^-/-^* P10 cerebella. Each dot represents the average of at least 3 images per animal. Data are mean ± s.d. *P*-values were determined by a two-tailed Student’s *t*-test.

## Discussion

Here, we identified a role for KIF3C in the developing mouse cerebellum. We found that *Kif3c* deletion results in reduced cerebellar size due to decreased CGNP proliferation. Unlike *Kif17* mutants and *Kif3a* mutants, *Kif3*c mutant cerebella do not exhibit altered HH signaling. In addition to decreased CGNP proliferation, we observe abnormal Bergmann glia density and localization and reduced expression of Notch target gene, *Hes1*.

### Microtubule Stability in Kif3c mutants

*Kif3c^-/-^* cerebella display decreased CGNP proliferation and disrupted Bergmann glia patterning. One potential function for KIF3C in cerebellar development is the regulation of microtubule stability. Previous work found *Kif3c^-/-^* neurons displayed impaired regeneration due to stable, overgrown, and looped microtubules at the growth cones of dorsal root ganglion neurons(Gumy et al., 2013). Further, KIF3C localized to the growing ends of microtubules and preferentially bound to tyrosinated (unstable) microtubules (Gumy et al., 2013). Forced homodimerization of KIF3C motors was also found to increase microtubule catastrophe frequency (Guzik-Lendrum et al., 2017).

Microtubules are essential in neuronal development. Cell migration, cue-dependent navigation of the growing end of axons and the arborization of dendrites all depend on microtubules [reviewed in (Baas et al., 2016)]. Bergmann glia project foot processes that attach to the basement membrane, which support the cerebellar cytoarchitecture. Purkinje cell axon growth and dendrite arborization are dependent on proper microtubule growth; CGNPs undergo vast remodeling of their cytoskeleton during differentiation and migration [reviewed in (Leto et al., 2016)]. It will be important to investigate the if KIF3C localizes to the tips of the growing ends of microtubules in the developing cerebellum. Further, examining *Kif3c^-/-^* cerebellar neurons to determine if they display abnormalities in their cytoskeleton or overgrown/looped microtubules will be vital to determining the contribution of *Kif3c* in cerebellar development.

### Redundancy and Compensation in Kinesin-2 Motors

The kinesin-2 family of motors consists of KIF3A, KIF3B, KIF3C and KIF17. Deletion of *Kif3a*, *Kif17,* or *Kif3c* all result in cerebellar hypoplasia (Spassky et al., 2008; Waas et al., 2024). Interestingly, while the deletion of either *Kif3a* or *Kif17* result in HH-dependent phenotypes, deletion of *Kif3c* does not perturb the levels of HH signaling.

The requirement of kinesin-2 motors varies significantly across different model organisms. For example, in *C. elegans*, the KIF3A/KIF3B homologue, KLP20/KLP11, contributes to building the middle segment of amphid-channel sensory cilia, while the KIF17 homologue, OSM-3, is responsible for generating the distal segment of cilia (Evans et al., 2006; Snow et al., 2004). However, OSM-3 can compensate for the loss of KLP20/KLP11, while KLP20/KLP11 cannot compensate in OSM-3 mutants. In the developing zebrafish, loss of *Kif17* results in ciliary defects in the retina and olfactory pit (Insinna et al., 2008; Lewis et al., 2018; Lewis et al., 2017; Zhao et al., 2012), suggesting the other kinesin-2 motors cannot compensate for its loss in those tissues. Loss of *Kif3b* in zebrafish results in delayed outer segment development and shortened cilia in the retina, suggesting *Kif17* or *Kif3c* can partially compensate for *Kif3b*. Further supporting this notion, injection of either *Kif17* or *Kif3c* mRNA in *Kif3b* mutant embryos can rescue ciliogenesis in specialized cell types (Zhao et al., 2012). However, this redundancy appears to be abolished in mice. Loss of *Kif3a* or *Kif3b* result in defective ciliogenesis and mid-gestation lethality (Nonaka et al., 1998; Takeda et al., 1999), while loss of *Kif17* or *Kif3c* does not result in ciliary phenotypes or embryonic lethality (Yang et al., 2001; Yin et al., 2011). Due to the lack of expression of *Kif3c* or *Kif17* during mouse embryogenesis, it is not surprising that these motors cannot compensate for KIF3A/KIF3B. However, overexpression of KIF17 or KIF3C in *Kif3a^-/-^;Kif3b^-/-^* NIH/3T3 cells cannot rescue ciliogenesis (Engelke et al., 2019), demonstrating expression of accessory kinesin-2 motors is not sufficient for compensation of KIF3A/KIF3B in mice. The cerebellum is an ideal tissue to examine the contribution of individual kinesin-2 motors because all motors are expressed during development and homeostasis.

While KIF3C loss did not impact HH signaling, it will still be interesting to examine kinesin-2 compound mutants. Examination of a true kinesin-2 null cerebellum (*En2Cre;Kif3a^fl/fl^;Kif17^-/-^*) will shed light on the requirement of the kinesin-2 family in cerebellar development (it is important to note that *Kif3a* deletion will be sufficient, as KIF3B/KIF3C has not been observed endogenously). Examination of mice with deletion of kinesin-2 motors specifically in Purkinje cells (*Shh^Cre/+^;Kif3a^fl/fl^;Kif17^fl/fl^*) or CGNPs (*Atoh1Cre;Kif3a^fl/fl^;Kif17^fl/fl^*) will also be crucial, considering two these cell types express all kinesin-2 motors. Kinesin-2 null Purkinje cells will be of particular importance, considering new studies demonstrating KIF3B function in SHH-producing cells in the developing limb (Wang et al., 2022), as well as the requirement of cilia in the survival of Purkinje cells (Bowie and Goetz, 2020).

## Supporting information

Supplement_KIF3C

## Acknowledgements

We thank past and present Allen lab members for their valuable feedback and suggestions. We thank members of the Department of Cell and Developmental Biology who provided access to equipment, including the O’Shea, Engel, and Spence labs. PAX6 and LIM1/2 antibodies were obtained from the Developmental Studies Hybridoma Bank, created by the Eunice Kennedy Shriver National Institute of Child Health and Human Development of the National Institutes of Health and maintained at The University of Iowa, Department of Biology, Iowa City, IA 52242, USA. Finally, we acknowledge the Biomedical Research Core Facilities Microscopy Core, which is supported by the Rogel Cancer Center, for providing access to confocal microscopy equipment.

## Notes

### Competing Interest Statement

The authors have declared no competing interest.

