## Supplement_KIF3C for "KIF3C Regulates Bergmann Glia Density and Patterning during Cerebellar Development"

**Supplemental Figures**

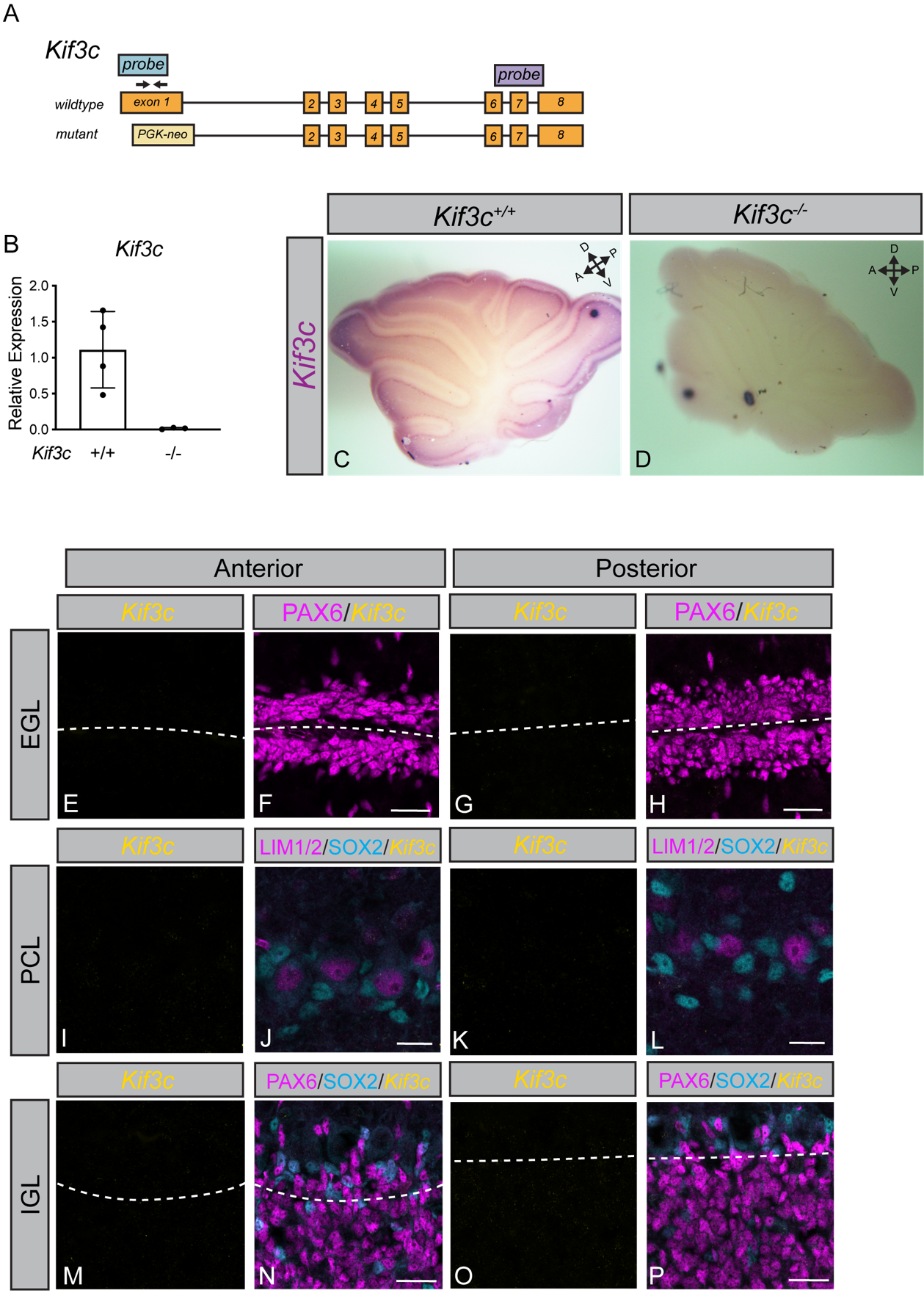

**Figure S1: *Kif3c* is expressed widely across the cerebellum.** Schematic of *Kif3c* wildtype and mutant alleles (**A**). The wildtype allele contains all 8 exons, while the mutant allele has insertion of a PGK-neo cassette immediately downstream of the start codon. Blue probe denotes the fluorescent *in situ* probe binding site, while the purple probe indicates the digoxigenin probe. Arrowheads above exon 1 represent RT-qPCR primers. RT-qPCR detection of *Kif3c* expression (**B**) in P10 *Kif3c^+/+^* and *Kif3c^-/-^* cerebella. Data are mean ± s.d. Each dot represents an individual animal. P-values were determined by a two-tailed Student’s t-test. Whole-mount *in situ* hybridization of *Kif3c^+/+^* (**C**) and *Kif3c^-/-^* (**D**) cerebella at postnatal day 10 (P10). Fluorescent *in situ* detection of *Kif3c* mRNA (yellow; **E-P**) in *Kif3c^-/-^* cerebella. Antibody detection of PAX6 (magenta) to label cerebellar granule neural progenitor nuclei in the EGL (**F, H**) Dashed lines (**E-H**) separate individual external granule layers. Antibody detection of LIM1/2 and SOX2 (magenta, cyan; **J, L**) to label Purkinje cells and Bergmann glia, respectively. Antibody detection of PAX6 and SOX2 (magenta, cyan, **N, P**) to label CGNs and Bergmann glia, respectively. Dashed lines (**M-P**) separate molecular layer (MCL) and internal granule layer (IGL). Scale bars (**F, H, N, P**), 25 μm. Scale bars (**J, L**), 15 μm.

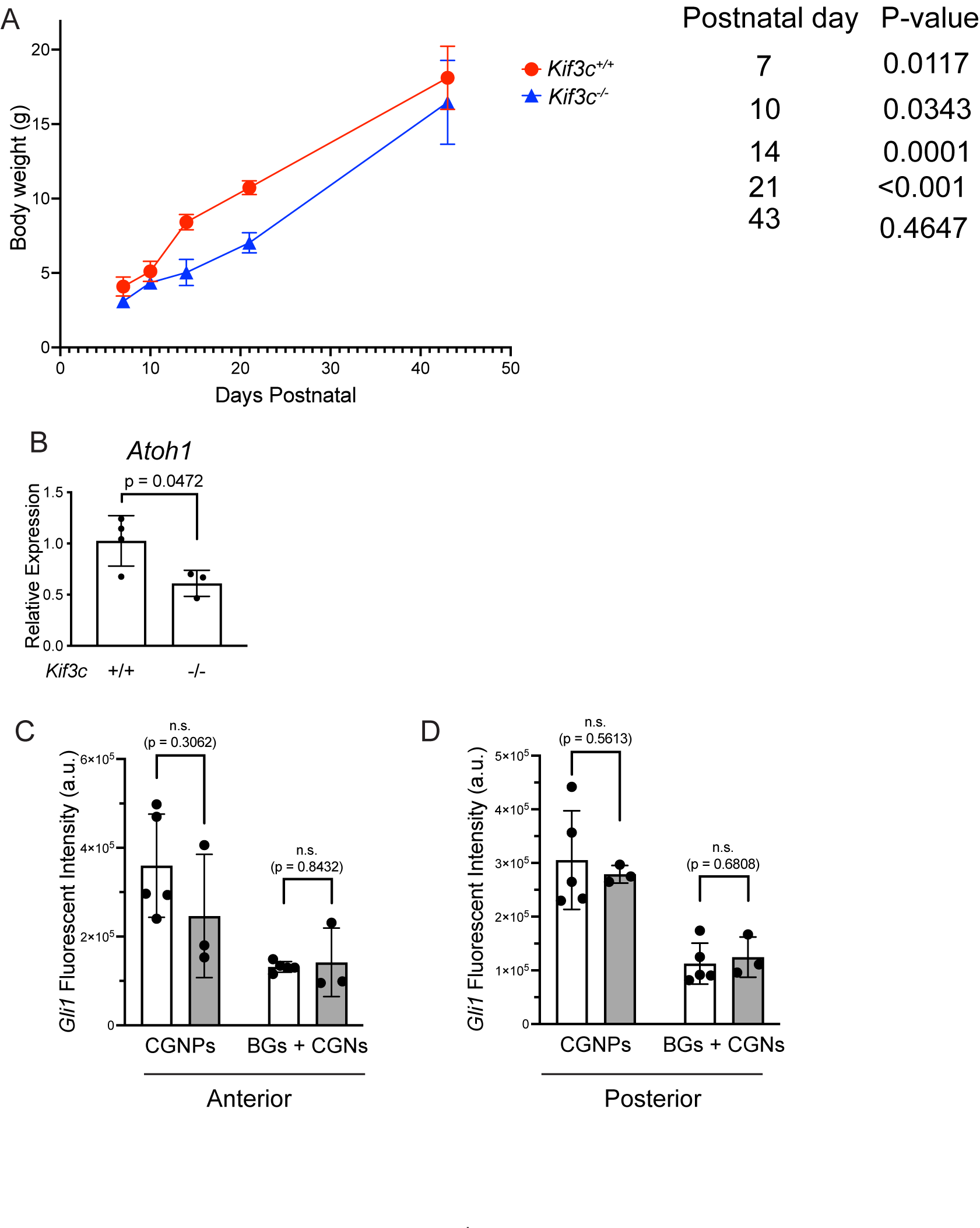

**Figure S2: *Kif3c* mutant mice remain smaller until adulthood.**

Mouse body weight over time (**A**) of *Kif3c^+/+^* and *Kif3c^-/-^* mice beginning at postnatal day 7 to 43. Each dot represents the average of a timepoint, with brackets for standard deviation. RT-qPCR detection of *Atoh1* expression (**B**) in P10 *Kif3c^+/+^* and *Kif3c^-/-^* cerebella. Quantitation of fluorescent intensity (**C, D**; integrated density) of *Gli1* puncta surrounding CGNPs and Bergmann glia (BG) and CGNs in the anterior (**C**) and posterior (**D**) lobes. Data are mean ± s.d. Each dot represents an individual animal. *P*-values were determined by a two-tailed Student’s *t*-test.

**Table 1: qPCR Primers**

| Gene | forward primer (5-3) | reverse primer (5-3) | Reference |
| --- | --- | --- | --- |
| *Gapdh* | GTGGTGAAGCAGGCATCTGA | GCCATGTAGGCCATGAGGTC | [Han et al., 2017 (PLoS Biology)] |
| *Kif3c* | CAGGCCGACCTGTATGACG | GTCCCCTGCATGGTGTAGG | designed by BW |
| *Atoh1* | AGTCAATGAAGTTGTTTCCC | ACAGATACTCTTATCTGCCC | [Hor et al., 2021 (Journal of Neuroscience)] |
| *Ptch1* | GAAGCCACAGAAAACCCTGTC | GCCGCAAGCCTTCTCTAGG | [Han et al., 2017 (PLoS Biology)] |
| *Ptch2* | CCCGTGGTAATCCTCGTGGCCTCTAT | TCCATCAGTCACAGGGGCAAAGGTC | [Shimokawa et al., 2008 (JBC)] |
| *Shh* | GCTGTGGAAGCAGGTTTCG | GGAAGGTGAGGAAGTCGCTC | [Madison et al., 2005 (Development)] |
| *Hes1* | CAGCCAGTGTCAACACGACAC | TCGTT CATGCACTCGCTGAAG | [Solecki et al., 2001 (Neuron)] |
| *Jag1* | TGCTTGGTGACAGCCTTCTACTGG | CTCTGGGCACTTTCCAAGTC | [Solecki et al., 2001 (Neuron)] |

**Table 2: Antibodies**

| Antibody | Source | Catalogue Number | Application | Concentration used |
| --- | --- | --- | --- | --- |
| Mouse IgG1 anti PAX6 | DSHB | PAX6 | IF | 1:20 |
| Mouse IgG1 anti LIM1+2 | DSHB | 4F2 | IF | 1:20 |
| Rabbit anti SOX2 | Seven Hills Bioreagent | WRAB-1236 | IF | 1:2000 |
| Rabbit anti Ki67 | Abcam | ab15580 | IF | 1:1000 |
| AP-conjugated anti-DIG antibody | Roche (Millipore Sigma) | 11093274910 | WISH | 1:4000 |
| Alexa Fluor 488 goat anti-mouse IgG1 | Invitrogen | A21121 | IF | 1:500 |
| Alexa Fluor 647 goat anti-mouse IgG1 | Invitrogen | A21240 | IF | 1:500 |
| Alexa Fluor 647 donkey anti-rabbit IgG | Invitrogen | A31573 | IF | 1:500 |
